# A method for allocating low-coverage sequencing resources by targeting haplotypes rather than individuals

**DOI:** 10.1101/188896

**Authors:** Roger Ros-Freixedes, Serap Gonen, Gregor Gorjanc, John M Hickey

## Abstract

**Background:** This paper describes a heuristic method for allocating low-coverage sequencing resources by targeting haplotypes rather than individuals. Low-coverage sequencing assembles high-coverage sequence information for every individual by accumulating data from the genome segments that they share with many other individuals into consensus haplotypes. Deriving the consensus haplotypes accurately is critical for achieving a high phasing and imputation accuracy. In order to enable accurate phasing and imputation of sequence information for the whole population we allocate the available sequencing resources among individuals with existing phased genomic data by targeting the sequencing coverage of their haplotypes.

**Results:** Our method, called AlphaSeqOpt, prioritizes haplotypes using a score function that is based on the frequency of the haplotypes in the sequencing set relative to the target coverage. AlphaSeqOpt has two steps: (1) selection of an initial set of individuals by iteratively choosing the individuals that have the maximum score conditional to the current set, and (2) refinement of the set through several rounds of exchanges of individuals. AlphaSeqOpt is very effective for distributing a fixed amount of sequencing resources evenly across haplotypes, which results in a reduction of the proportion of haplotypes that are sequenced below the target coverage. AlphaSeqOpt can provide a greater proportion of haplotypes sequenced at the target coverage by sequencing less individuals, as compared with other methods that use a score function based on the haplotypes population frequency. A refinement of the initially selected set can provide a larger more diverse set with more unique individuals, which is beneficial in the context of low-coverage sequencing. We extend the method with an approach to filter rare haplotypes based on their flanking haplotypes, so that only those that are likely to derive from a recombination event are targeted.

**Conclusions:** We present a method for allocating sequencing resources so that a greater proportion of haplotypes are sequenced at a coverage that is sufficiently high for population-based imputation with low-coverage sequencing. The haplotype score function, the refinement step, and the new approach of filtering rare haplotypes make AlphaSeqOpt more effective for that purpose than methods reported previously for reducing sequencing redundancy.

## Introduction

This paper describes a heuristic method for allocating low-coverage sequencing resources by targeting haplotypes rather than individuals so that haplotypes have a coverage that is sufficiently high for population-based imputation.

The use of whole-genome sequencing data has great potential in livestock breeding programs. It may increase the power of discovery of causative variants [1–3] and may enable more accurate and persistent predictions of breeding values than marker array genotypes [4,5]. To capture the full potential of sequence data in livestock, sequence and phenotype data on a large number, perhaps millions, of individuals may be required to accurately estimate the effects of the large number of causative variants that underlie quantitative traits [6].

Low-cost sequencing strategies combined with imputation can be utilised to generate the required amount of sequence information for a large number of individuals at an affordable cost [7–11]. The strategies for low-cost sequencing can be classified into three groups: (1) to sequence a certain number of key individuals at high coverage, as in the 1,000 Bull Genomes project (**KeySires**) [2,5]; (2) to sequence a larger number of individuals at low coverage (**LCSeq**) [6,12,13]; and (3) to sequence a set of chosen individuals at a wide range of coverages (**VarCoverage**) [14].

The LCSeq approach exploits the fact that the population structures that are typical in livestock breeding result in individuals being sufficiently related to share large genome segments. LCSeq focuses sequencing on the haplotypes in the population rather than on any individual. LCSeq sequences individuals at low coverage and assembles high-coverage sequence information for every haplotype by accumulating the low-coverage sequence data from the genome segments that are shared between many individuals to derive the ‘consensus haplotypes’. The consensus haplotypes are then used to impute the sequence data of the individuals. Deriving the consensus haplotypes accurately is critical for achieving a high phasing and imputation accuracy under the LCSeq strategy.

With the LCSeq approach potentially many more individuals can be sequenced than with the KeySires or VarCoverage approaches. This provides three advantages to the LCSeq approach: (1) higher variant discovery rates, particularly for low-frequency variants [15]; (2) inclusion of rare haplotypes; and (3) a more precise capture of the recombination events that have occurred in the population, which would enable better definition of the haplotypes that are present in the population and thus better imputation of these haplotypes into the individuals that carry them.

There are methods to optimise the selection of individuals for sequencing for the three alternate sequencing approaches. Most of these methods focus only on the choice of which individuals to sequence with the aim to impute their sequence information into their relatives [5,16–18]. Recently, Gonen et al. [14] proposed a method that identifies the individuals with the largest genetic footprint on the population and optimises the allocation of sequence resources across these focal individuals and their ancestors with the aim to maximise phasing accuracy of their sequenced haplotypes when using family-based phasing methods.

Although LCSeq could be used alone, we envisage a sequencing strategy in two stages for facilitating the imputation of sequence data. The first stage uses the method developed by Gonen et al. [14] with the aim of producing a set of accurately phased haplotypes that are shared by a lot of individuals in the population. The second stage seeks to complement the first stage by applying the LCSeq approach as described above to spread low-coverage sequence data across the population so that whole-genome sequence data can be imputed to the whole population, which in turn will be enhanced by the phasing of the most common haplotypes achieved in the first stage. To do this effectively a method for optimising the allocation of sequencing resources under the LCSeq approach should be developed.

We hypothesise that such a method should maximise the sequencing coverage of the maximum possible number of haplotypes because this would enable population-based phasing and imputation methods rather than family-based imputation methods to accurately phase and impute the data to all individuals. For such population-based phasing and imputation methods, a certain level of sequence coverage must be accumulated for accurate inference of a consensus haplotype. With a prototype of such a population-based phasing and imputation method we observed that there is a positive relationship between the coverage that a particular haplotype accumulates across individuals and the imputation accuracy of a consensus haplotype (for a description, see Additional file 1: Figure S1). A random allocation of sequencing resources under the LCSeq approach results in some haplotypes being sequenced many times, some rarely, and some not at all. To optimise the allocation of sequencing resources under LCSeq we need to maximise the proportion of haplotypes that are sequenced at the target coverage and minimise the proportion of haplotypes that are under-or over-sequenced. Similarly, we need to minimise the sequencing resources allocated to haplotypes that are too rare to have consensus haplotypes inferred or their effects estimated accurately.

The objective of this work was to develop a method that uses haplotypes derived from existing phased marker array genotypes to identify which individuals should be sequenced, and at what coverage, to maximize the proportion of consensus haplotypes sequenced at a minimum target coverage. Our method uses a score function to identify a set of individuals based on the coverage at which their haplotypes are sequenced and then it refines the initial set of individuals through rounds of exchanges. We extend the method with an approach to filter out rare haplotypes so that we only target those that are likely to derive from the recombination of common haplotypes. We tested the performance of the algorithm using simulated data and the results showed that our method is efficient in distributing the sequencing resources evenly across a large proportion of the haplotypes observed in the population.

## Materials and Methods

### Description of the AlphaSeqOpt method

Our method utilises existing phased marker array genotypes to identify which individuals should be sequenced, and at what coverage, so that the maximum proportion of haplotypes are sequenced at any minimum target coverage with a fixed sequencing budget. The method has two main steps. In the first step, referred as ‘initial set selection’, an initial set of individuals is selected by iteratively choosing the individuals that are the most complementary to the ones already in the set according to a score function. In the second step, referred as ‘set refinement’, the initial set of individuals is refined through several rounds of exchanges. The method was implemented in a software package called AlphaSeqOpt, which also implements the method of Gonen et al [14]. Throughout the rest of the paper, AlphaSeqOpt is used when referring to our method.

#### Initialisation step

0a: Construct a haplotype library for the population using existing phased marker array genotypes. Split each chromosome into *c* cores of length *m* markers. A ‘core’ is each of the strings of *m* consecutive marker positions used to determine the haplotypes. Within a core, strings of alleles (previously phased) are compared to define which haplotype each individual carries in each parental chromosome. Strings of alleles that are identical between two individuals are defined as a unique haplotype and strings with multiple mismatches are defined as different haplotypes. A predefined number of mismatches can be allowed before two strings of alleles are defined as different haplotypes to account for sequencing errors.

0b: Calculate the maximum size of the sequencing set. Assuming linear sequencing costs, the sequencing budget divided by the cost of 1x sequencing determines the total amount of sequencing coverage that could be produced, represented by the number of slots of the sequencing set. A ‘slot’ is each of the positions in the set, which can be assigned to any given individual following the steps below. Each slot corresponds to 1x sequencing.

#### Initial set selection (step 1)

1a: Calculate a score for each haplotype in each core. We derived a score function that prioritizes the haplotypes that are closer to reaching the target coverage. The score function is based on the frequency of a haplotype in the sequencing set relative to the target coverage. The score function is:

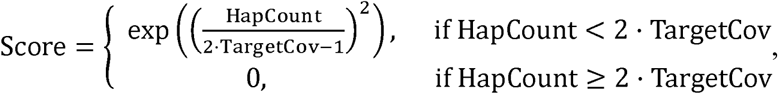

where HapCount is the number of times that a haplotype appears in the current sequencing set and TargetCov is the target haplotype coverage (Figure 1). The score increases every time that an individual that carries a given haplotype is added to the sequencing set. When the haplotype count in the set reaches twice the target coverage, which is the haplotype count required to produce the target coverage assuming that for each x of coverage of an individual there is a probability of 0.5 of reading either the paternal or maternal haplotype, the score is set to 0 to prevent over-sequencing of well-covered haplotypes in favour of allocating sequencing resources to other haplotypes.

**Figure 1.**
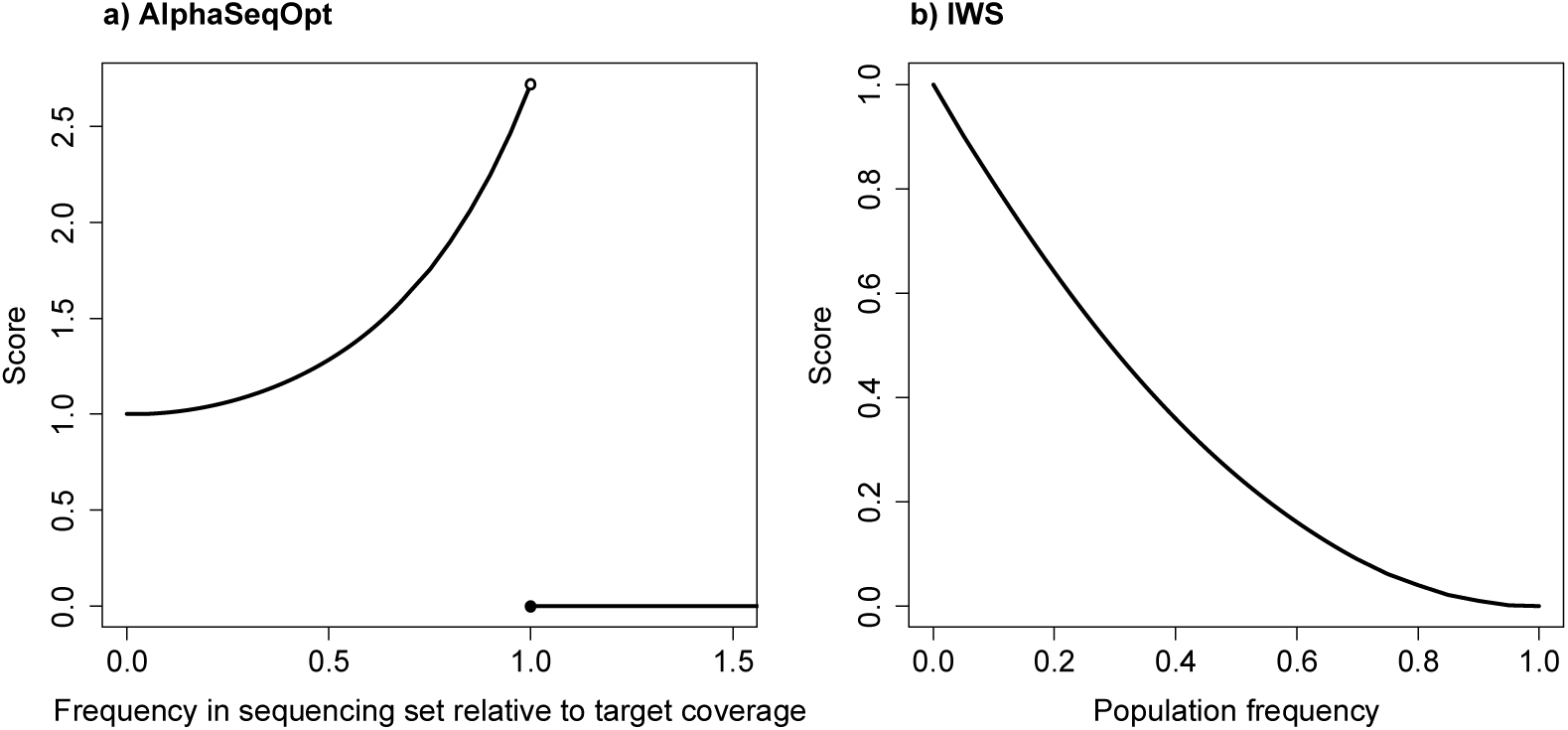
Score functions: (a) in AlphaSeqOpt; and (b) in the IWS method proposed by Bickhart et al. [17]. Note the different axes: in (a) scores range from 1 to e^*k*^ based on the frequency of the haplotype in the sequencing set relative to the target coverage, which is variable across rounds; in (b), scores range from 0 to 1 based on the population frequency, which is fixed.

1b: Calculate the total score for every individual as the sum of the scores of the haplotypes that each individual carries at each core.

1c: Add the individual with the maximum score to the first available slot of the initial set. If there is more than one individual satisfying this condition, one individual is selected at random amongst those individuals with the maximum score. Repetition of individuals in several slots of the set is allowed. The number of slots occupied by the same individual indicates at what coverage it should be sequenced (i.e., an individual that appears *n* times in the set should be sequenced at *n*x).

1d: Calculate the total cost of sequencing the current set as the cost of library preparation times the number of individuals in the set plus the cost of 1x sequencing times the total sequencing coverage produced.

1e: Repeat steps 1a to 1d until the initial set is complete (i.e., we have a set of individuals at variable coverage that exhausts all the sequencing resources). Because some resources are used for library preparation some slots will be left empty.

#### Set refinement (step 2)

2a: Choose randomly a predetermined number of slots of the set. Remove the individuals assigned to these slots from the set.

2b: Repeat steps 1a to 1d to fill the emptied slots. Individuals removed in step 2a can go back into the set if they have the maximum individual score.

2c: If the exchanges result in the same or a greater percentage of unique haplotypes sequenced at (or above) the target coverage, keep the new set. Otherwise, discard the new set in favour of the previous set.

2d: Repeat steps 2a to 2c for a predefined number of exchange rounds.

If there are individuals that have been sequenced previously, AlphaSeqOpt can account for the available sequence data easily by adding the pre-existing coverage of their haplotypes to HapCount during the calculation of the haplotype scores in step 1a. If there are not any individuals that have been sequenced previously, all haplotypes will have the same starting score and the first individual will be selected at random amongst those that have more non-missing haplotypes.

For any given target haplotype coverage, AlphaSeqOpt will produce a set of individuals to be sequenced from 1x to a maximum coverage equal to twice the target coverage. To ensure that all individuals are sequenced at a low coverage and that a larger number of individuals is sequenced it is also possible to restrict the coverage for the individuals in the set to a desired maximum (e.g., to 1x or 2x).

In the implementation of AlphaSeqOpt we are making two assumptions regarding the yield of data from the sequencer: (1) that sequencing coverage is uniform across the genome; and (2) that for each x of coverage of an individual there is a probability of 0.5 of reading either the paternal or maternal haplotype and therefore each haplotype receives half the coverage. Even though these assumptions contradict empirical observations [19], there is no straightforward way of accounting for variation of coverage across genome or between alleles prior to performing sequencing. Regarding the sequencing costs, we are assuming that when we increase the sequencing coverage we incur a linear increase of the sequencing costs. AlphaSeqOpt can also account for non-linear cost structures by modifying the cost equation used in step 1d.

### Algorithm testing

The proposed method was tested against our implementation of the Inverse Weight Selection (IWS) method as described by Bickhart et al. [17], our adaptation of the IWS method to obtain more comparable results, and a method that selects the individuals randomly (referred to as Random).

The IWS method as described by Bickhart et al. [17] follows the step 1 as described above but in step 1a it uses an inverted parabolic score function 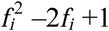, where *ƒ*_*i*_ is the population frequency of the haplotype. Note that this function uses the population frequency, while the score function that we propose uses the frequency of the haplotype in the sequencing set relative to the target coverage. The two score functions are compared in Figure 1. Another major difference with AlphaSeqOpt is that Bickhart et al. [17] proposed targeting only homozygous haplotype cores based on the marker array genotypes. Thus, the IWS method only scores such haplotypes and it stops after the initial set is constructed, without a step of refinement.

Our adaptation of IWS mirrored the method that we propose more closely, including a step of refinement of the initially selected set, with the only difference being the score function used. This method follows both steps 1 and 2 as described above but in step 1a it uses the inverted parabolic function 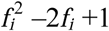. We did not follow the suggestion of targeting only the haplotypes at cores that are predicted to be homozygous based on the marker array genotypes, because this would disadvantage the adapted IWS method.

The Random method also used the algorithm described but individuals were selected randomly instead of according to a score function. In the refinement step, random exchanges of individuals were performed.

All methods were tested in a range of scenarios. The scenarios varied in the target haplotype coverage (5x, 10x, or 15x) and in the total available sequencing resources (£400,000, £800,000, or £1,600,000 GBP). We calculated the cost of each scenario assuming a cost in library preparation of £40 and a cost in 1x sequencing of £80. The tested sequencing resources would produce a total of 5,000x, 10,000x, or 20,000x whole-genome reads, respectively, if cost of library preparation was ignored. Haplotypes observed only once or twice in the population were excluded from the analyses unless stated otherwise. Additional tests were performed with a restriction of maximum individual coverage of 1x, for different numbers of exchanges per round, ranging from 1 slot to the total size of the set, and for different costs of library preparation, ranging from no cost to £40. We performed 10 repetitions for all analyses. The percentage of unique haplotypes sequenced at (or above) the target coverage was used as the main criterion, together with the number of individuals sequenced.

For simplicity, in some instances we will focus on the scenarios with a target haplotype coverage of 10x but the algorithm can be used with any desired target coverage.

### Filtering of rare haplotypes based on flanking context

A new approach for filtering the rare haplotypes included in the analyses was also developed. In this approach we filtered the rare haplotypes so that only those rare haplotypes that are likely to derive from a recombination event between two common haplotypes were targeted.

The filtering was based on two assumptions: (1) rare haplotypes that were derived from a recombination event between common haplotypes will be flanked by common haplotypes; and (2) there will be no other individuals that carry the same combination of haplotypes at the cores that flank the rare recombined haplotype. The second assumption could be false if, for example, there had been multiple recombination events at different positions of the same core that produced multiple rare recombinant haplotypes from the same two common haplotypes, but note that this is a method for directing the sequencing resources among rare haplotypes, not an exact method for capturing all recombination events. Note also that combinations of consecutive cores with rare haplotypes could indicate either genomes that are unrelated to the population or phasing errors.

We implemented the above filtering approach according to the population count of the haplotypes at each core. In any given core, haplotypes with population count below a predefined threshold are included in the analysis only if all of the following conditions are met: (1) the rare haplotype is not at the first or last core of a chromosome; (2) the counts of the flanking haplotypes are greater than a predefined threshold (FlankCount); and (3) there are less than a predefined number (nComb) of individuals carrying the same combination of haplotypes flanking the rare haplotype.

In our implementation of AlphaSeqOpt we used this filtering approach on those rare haplotypes with population count ≤2 (observed only once in the population, referred to as ‘singletons’, or twice, referred to as ‘doubletons’) using FlankCount=2 and nComb=3. The same method could be applied for any population count. This approach for filtering the rare haplotypes was tested against the reference case with no filtering and against the approach in which all singletons and doubletons were filtered out.

### Simulated dataset

To demonstrate the implementation of the algorithm, a testing dataset was simulated to mimic a typical livestock population with known structured pedigree.

Sequence data was generated for 1,000 base haplotypes for each of ten chromosomes using the Markovian Coalescent Simulator [20] and AlphaSim [21,22]. Chromosomes were simulated to be 100 cM and 10^8^ base pairs in length, with a per site mutation rate of 2.5×10^−8^ and a per site recombination rate of 1.0×10^−8^. The effective population size (N_e_) was set to specific values during the simulation based on previously estimated N_e_ values within the Holstein cattle population [23]. These set values were: 100 in the base generation, 1,256 at 1,000 years ago, 4,350 at 10,000 years ago, and 43,500 at 100,000 years ago, with linear changes in between. The resulting sequence had approximately 650,000 segregating SNP loci across the ten chromosomes.

To enable the selection of sires for the generation of a pedigree, a quantitative trait influenced by 10,000 QTN distributed equally across the ten chromosomes was simulated. QTN positions were randomly chosen from the 650,000 segregating sequence loci and their effect sizes sampled from a normal distribution with a mean of zero and standard deviation of 0.01 (1.0 divided by the square root of the number of QTN). The QTN effects were used to compute the true breeding value (TBV) for each individual.

To emulate livestock breeding populations, a pedigree of 15 generations was simulated. Each generation comprised 1,000 individuals in equal sex ratio (i.e., 500 males and 500 females). In the first generation, chromosomes for each individual were sampled from the 1,000 sequence haplotypes in the base generation. In subsequent generations, chromosomes of each individual were sampled from parental chromosomes, assuming recombination with no interference. In each generation, the 25 males with the highest TBVs were selected as sires of the next generation. No selection was performed on females, and all 500 females were used as parents.

All individuals were assumed to be genotyped with a panel of 10,000 SNP markers distributed equally across the ten chromosomes. Marker genotypes of all individuals were phased using AlphaPhase [24–26] as input for AlphaSeqOpt. The parameters used for determining the population haplotype libraries were: (1) population haplotype libraries were created using individuals and SNPs with at least 90% phased genotype data; (2) sharing of haplotypes was determined as 100% identity matches; and (3) core lengths were set to 100 SNPs per chromosome.

In summary, the algorithm was tested using a dataset with 15,000 individuals. Individuals had 10 chromosomes and 10 cores per chromosome. The total number of haplotypes in the population was 8850 (on average, 88.5 haplotypes per core). Further details on the simulated dataset can be found in Gonen et al. [14].

### Software availability

The method has been implemented in the AlphaSeqOpt software package. AlphaSeqOpt is available for download at http://www.alphagenes.roslin.ed.ac.uk/alphaseqopt/, along with a detailed user manual.

## Results

### Performance of algorithm

AlphaSeqOpt allocated sequencing resources to enable a greater percentage of haplotypes in the population to be sequenced at the target coverage than other methods previously reported.

Figure 2 shows the comparison of AlphaSeqOpt with IWS, the adapted IWS, and Random when the target haplotype coverage was 10x. We tested different scenarios in which the total available sequencing resources were £400,000, £800,000, or £1,600,000. Figure 2a shows the percentage of haplotypes that would be sequenced at (or above) the target coverage of 10x by sequencing the set of individuals selected with AlphaSeqOpt. Figure 2b shows the number of individuals selected for sequencing in each of the scenarios. AlphaSeqOpt delivered the highest percentage of haplotypes sequenced at the target coverage, followed by the adapted IWS method, which achieved a lower percentage even though it sequenced a number of individuals similar to AlphaSeqOpt. The IWS method resulted in only a very small set of individuals being sequenced and these individuals captured only a small percentage of haplotypes sequenced at the target coverage. This result occurred because, as done by Bickhart et al. [17], we only targeted haplotypes that appeared in a homozygous state in at least one animal, which represent a small proportion of the haplotypes observed in the population, and therefore the IWS method did not exhaust all the available sequencing resources in any of the cases tested. The Random method sequenced a very large set of individuals but it was inefficient for obtaining the haplotypes sequenced at the target coverage.

**Figure 2.**
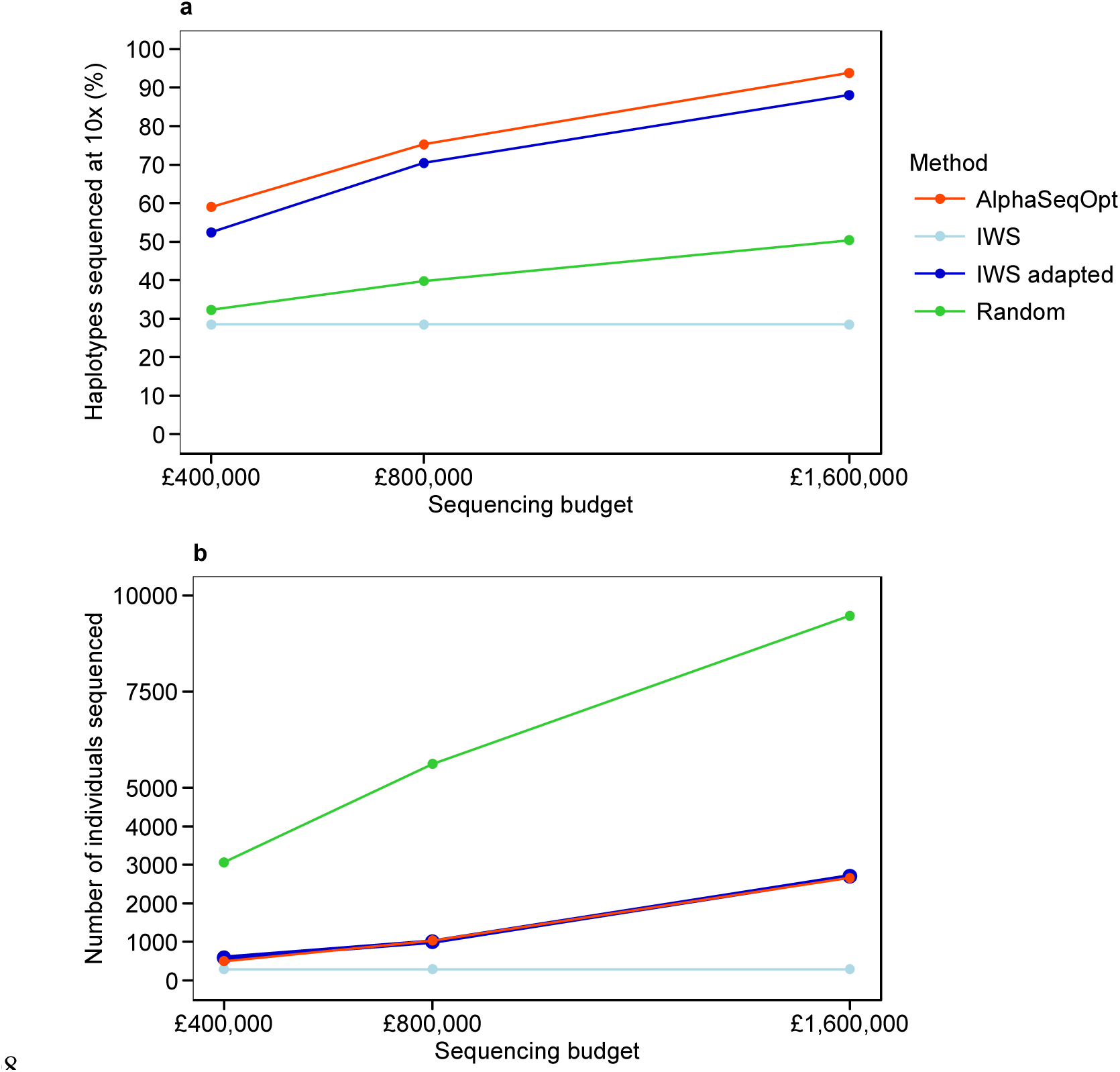
Performance of the four methods tested with different amounts of sequencing resources, in terms of: (a) the percentage of population haplotypes sequenced at (or above) the target coverage of 10x; and (b) the number of individuals sequenced. Standard errors were less than 0.2% (a) and 25 (b).

The AlphaSeqOpt method was further tested to assess the effect of its main features on the percentage of haplotypes sequenced at (or above) the target coverage, the number of individuals sequenced, the performance under restriction of the maximum coverage per individual, and the performance of the refinement step with different number of exchanges per round.

#### Percentage of haplotypes sequenced at the target coverage

The advantage provided by the AlphaSeqOpt score function and the step of refinement over the adapted IWS method is shown in Figure 3. Figure 3a shows the percentage of the haplotypes that would be sequenced at (or above) the target coverage by sequencing the set of individuals selected with AlphaSeqOpt. We tested nine scenarios in which the target coverage was 5x, 10x, or 15x and the total available sequencing resources were £400,000, £800,000, or £1,600,000. Each scenario was tested with either the AlphaSeqOpt score function or the IWS score function (adapted IWS method), and both the initial and refined sets were examined.

**Figure 3.**
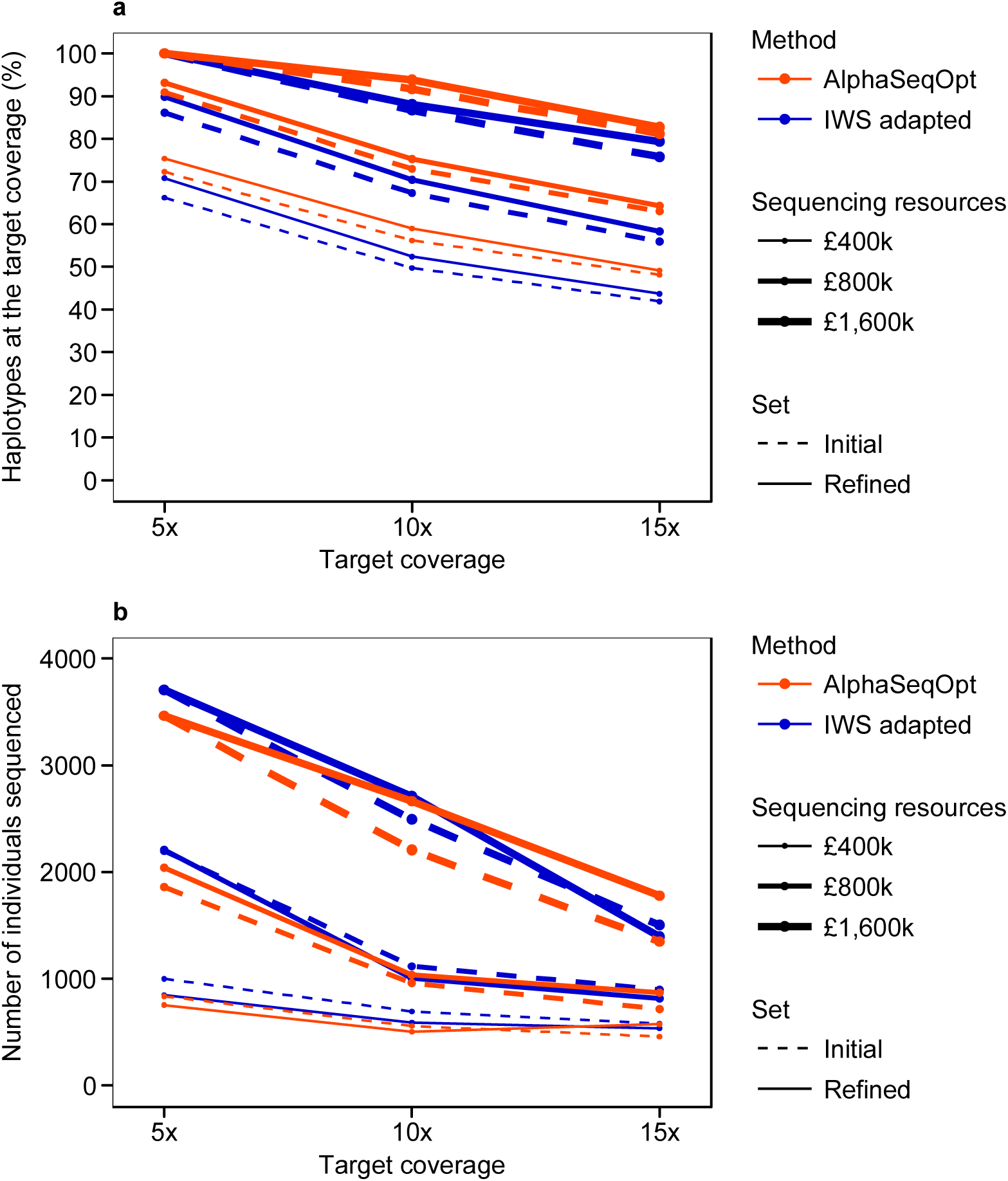
Performance of AlphaSeqOpt and the adapted IWS method with three levels of target haplotype coverage and different amounts of sequencing resources, in terms of: (a) the percentage of population haplotypes sequenced at (or above) the target coverage of 10x; and (b) the number of individuals sequenced. Both the initial and refined sets were examined. Standard errors were less than 0.4% (a) and 40 (b).

The AlphaSeqOpt score function provided a greater percentage of haplotypes sequenced at the target coverage than the IWS score function in all scenarios. The AlphaSeqOpt score function gave 1.8 to 6.6% more haplotypes sequenced at the target coverage than the IWS score function. The advantage of the AlphaSeqOpt score function was observed both in the initial and refined sets. The refinement step increased the percentage of haplotypes sequenced at the target coverage by 1.0 to 3.1% with the AlphaSeqOpt score function and 1.4% to 4.7% with the IWS score function. In total, using the AlphaSeqOpt score function and a refinement step delivered 6.6 to 9.3% more haplotypes sequenced at the target coverage than using the IWS score function without a refinement step.

AlphaSeqOpt performed better because it was more efficient at allocating the sequencing resources so that there were very few haplotypes that received some, but insufficient, sequencing coverage.

Figure 4a shows the distribution of the population count of the haplotypes and Figure 4b the distribution of the sequencing coverage that the haplotypes receive by sequencing the set of individuals selected with each method. Note that the x-axis in Figure 4b is half that of Figure 4a because for each x of coverage of an individual there is a probability of 0.5 of reading either the paternal or maternal haplotype in the diploid species that was simulated. Because the results for all scenarios were similar, for illustration purposes from here onwards we only show results for the scenario in which the target coverage was 10x and the sequencing resources were £800,000. Also note that the haplotypes with population count ≤2 are shown in Figure 4a but were excluded from the analyses shown in Figure 4b.

**Figure 4.**
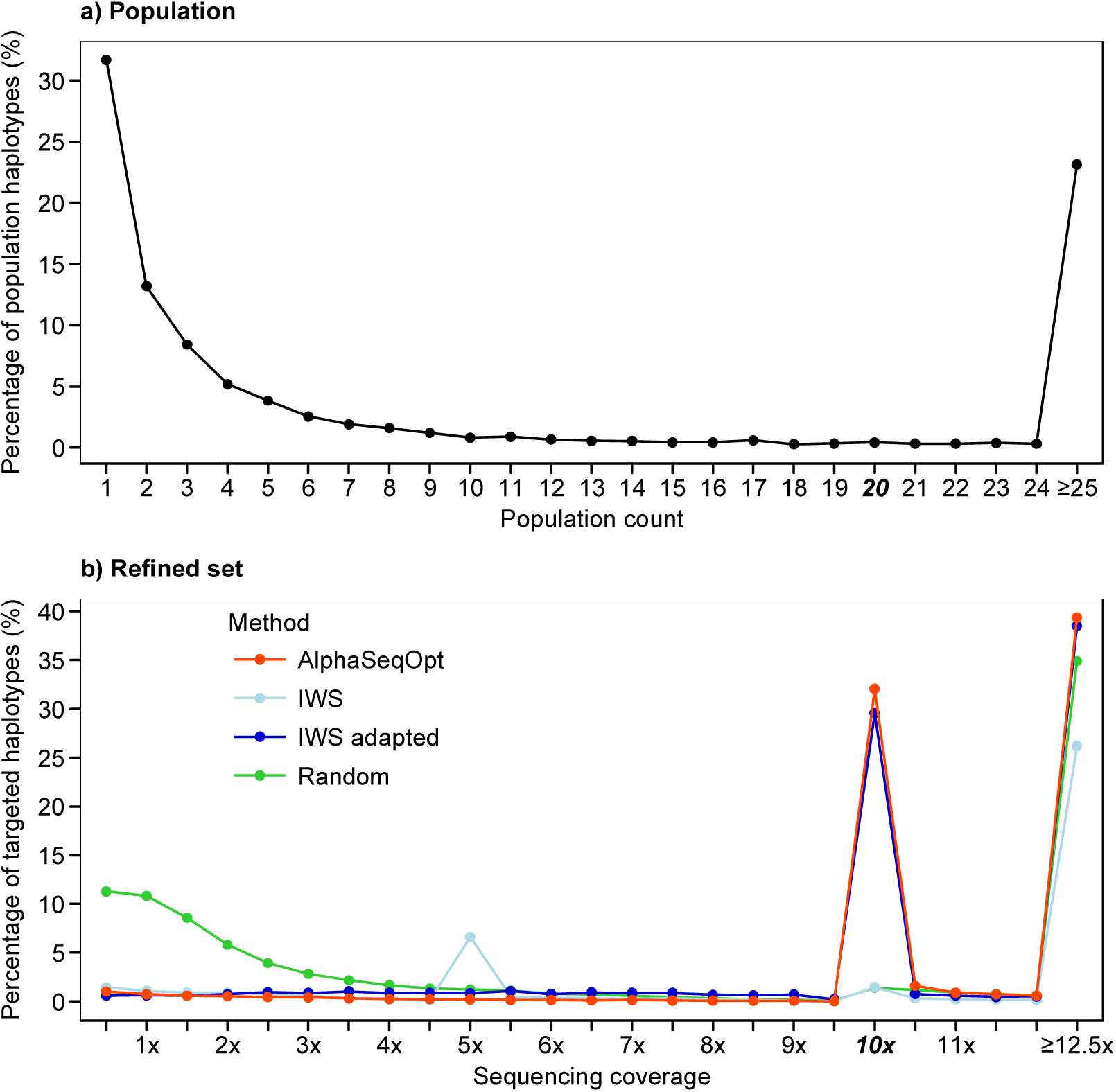
Distribution of: (a) the population count of the haplotypes; and (b) the sequencing coverage of the haplotypes using the four methods tested. The target haplotype coverage was 10x, the sequencing resources were set to £800,000 and haplotypes with population count ≤2 were excluded from the analyses. Standard errors were less than 14 (b).

As a reference, choosing individuals randomly followed by random exchanges of individuals followed the distribution of the population frequencies, with a large percentage of haplotypes sequenced at coverages below the target 10x (54.0% of the haplotypes had sequence coverage between 0.5x and 9.5x). The AlphaSeqOpt score function reduced this percentage to only 6.3% in the initial set and 5.6% in the refined set. This percentage was greater with the adapted IWS method than with AlphaSeqOpt in both sets (17.3% in the initial set was reduced to 14.7% in the refined set). The percentage of haplotypes that received no coverage at all in the refined set were 19.2% for AlphaSeqOpt, 14.9% for the adapted IWS, and 6.3% for Random.

#### Number of individuals sequenced

The initial sets that were selected by AlphaSeqOpt produced greater percentages of haplotypes at the target coverage by sequencing less animals than the sets selected by the adapter IWS method. The refinement step with the AlphaSeqOpt score function produced sequencing sets that contained a larger number of unique individuals than with the IWS score function. The extent to which the size of the sequencing set was increased depended on the cost of library preparation and the amount of sequencing resources available.

Figure 3b shows the number of individuals in the sets selected in each of the scenarios explored in Figure 3a. The initial set was smaller with the AlphaSeqOpt score function than with the IWS score function by between 122 and 340 individuals. During the refinement step with the AlphaSeqOpt score function, the set maintained approximately the same size when a small amount of sequencing resources was available but increased by up to 457 individuals when more sequencing resources were available. In contrast, during the refinement with the IWS score function, the size of the sequencing set decreased when few sequencing resources were available but remained more stable with a large amount of sequencing resources.

Figure 5 shows the effect of the cost of library preparation on the percentage of haplotypes sequenced at (or above) the target coverage (Figure 5a) and the number of unique individuals (Figure 5b) in the refined set produced with the AlphaSeqOpt score function or the IWS score function. With both score functions, the percentage of haplotypes sequenced at the target coverage increases linearly with decreasing library costs. When library cost is low, the AlphaSeqOpt score function produces larger sets with more unique individuals than the IWS score function, and these larger sets produce greater percentages of haplotypes sequenced at the target coverage. When the library costs are high, the difference between the sizes of the sets obtained with the two score functions is reduced. Figure 5c shows the distribution of the sequencing coverage across sequenced individuals in the refined set produced with the AlphaSeqOpt score function considering two extreme library costs. Low library costs allowed for the sequencing of more individuals at low coverage while high library costs resulted in a greater number of individuals being sequenced at twice the target coverage of the haplotypes. With a library cost of £5 the number of individuals sequenced was 1307.6 (302.2 at 1x to 124.3 at 20x) and with a library cost of £40 it decreased to 1036.4 (136.6 at 1x to 176.7 at 20x).

**Figure 5.**
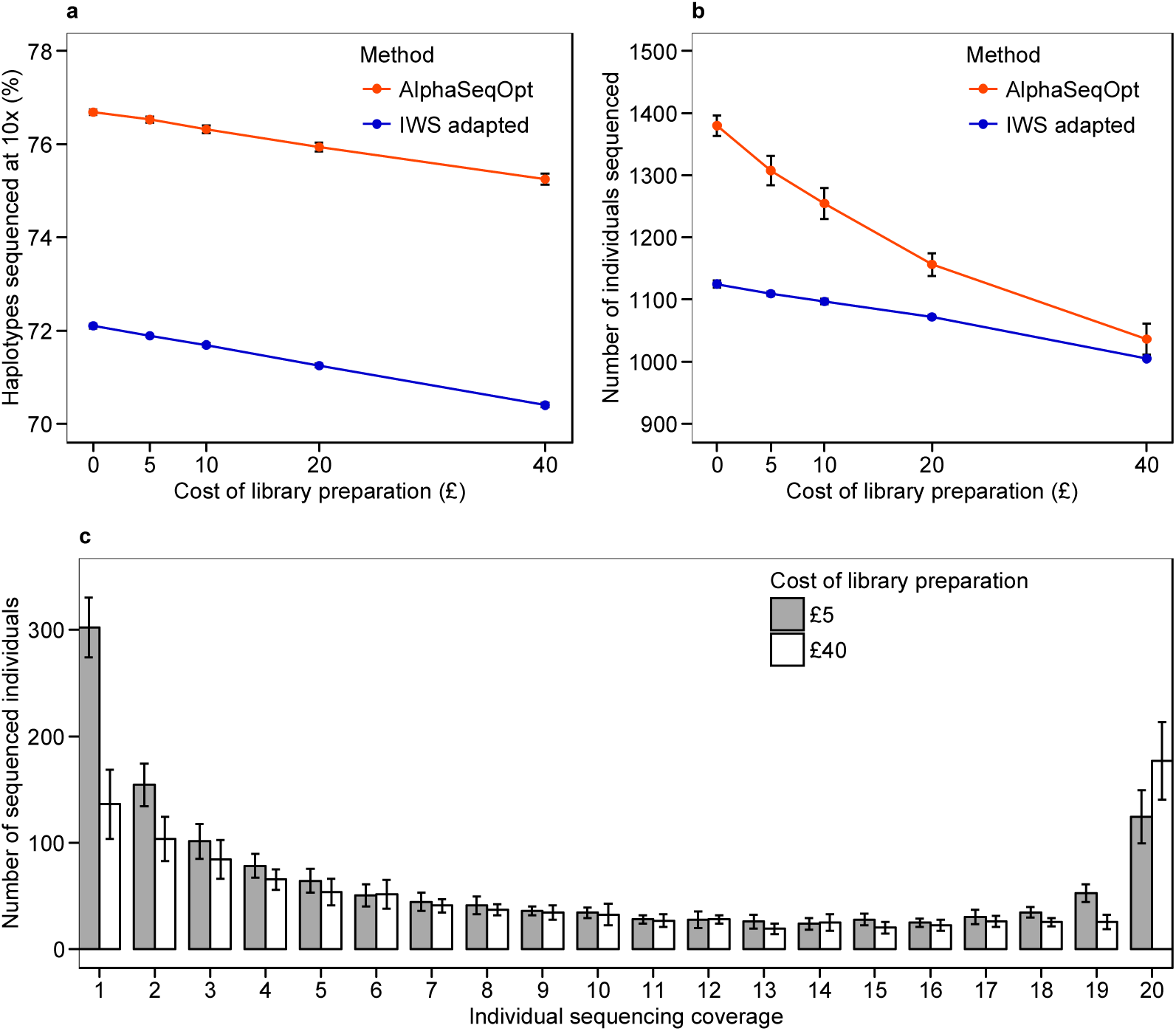
Effect of the cost of library preparation: (a) on the percentage of haplotypes sequenced at (or above) the target coverage of 10x; (b) on the number of individuals sequenced, using AlphaSeqOpt or the adapted IWS method; and (c) on the distribution of sequencing coverage of the individuals selected with AlphaSeqOpt. The sequencing resources were set to £800,000 and haplotypes with population count ≤2 were excluded from the analyses.

#### Restriction of individual coverage

The size of the sequencing set can be maximised by restricting the maximum coverage that each individual can get, so that the target coverage of the haplotypes is achieved by accumulating individuals sequenced only at or below a certain coverage. Figure 6 shows the comparison of AlphaSeqOpt with the adapted IWS when the maximum individual coverage is restricted to 1x. Under this restriction, only haplotypes with a population count ≥10, ≥20, and ≥30 can reach the target coverages of 5x, 10x, and 15x, respectively, and therefore haplotypes with lower population counts were excluded from the analyses. Figure 6a shows the percentage of targeted haplotypes that would be sequenced at (or above) the three levels of target coverage by sequencing the set of individuals selected with AlphaSeqOpt. Figure 6b shows the number of individuals selected for sequencing in each of the scenarios.

**Figure 6.**
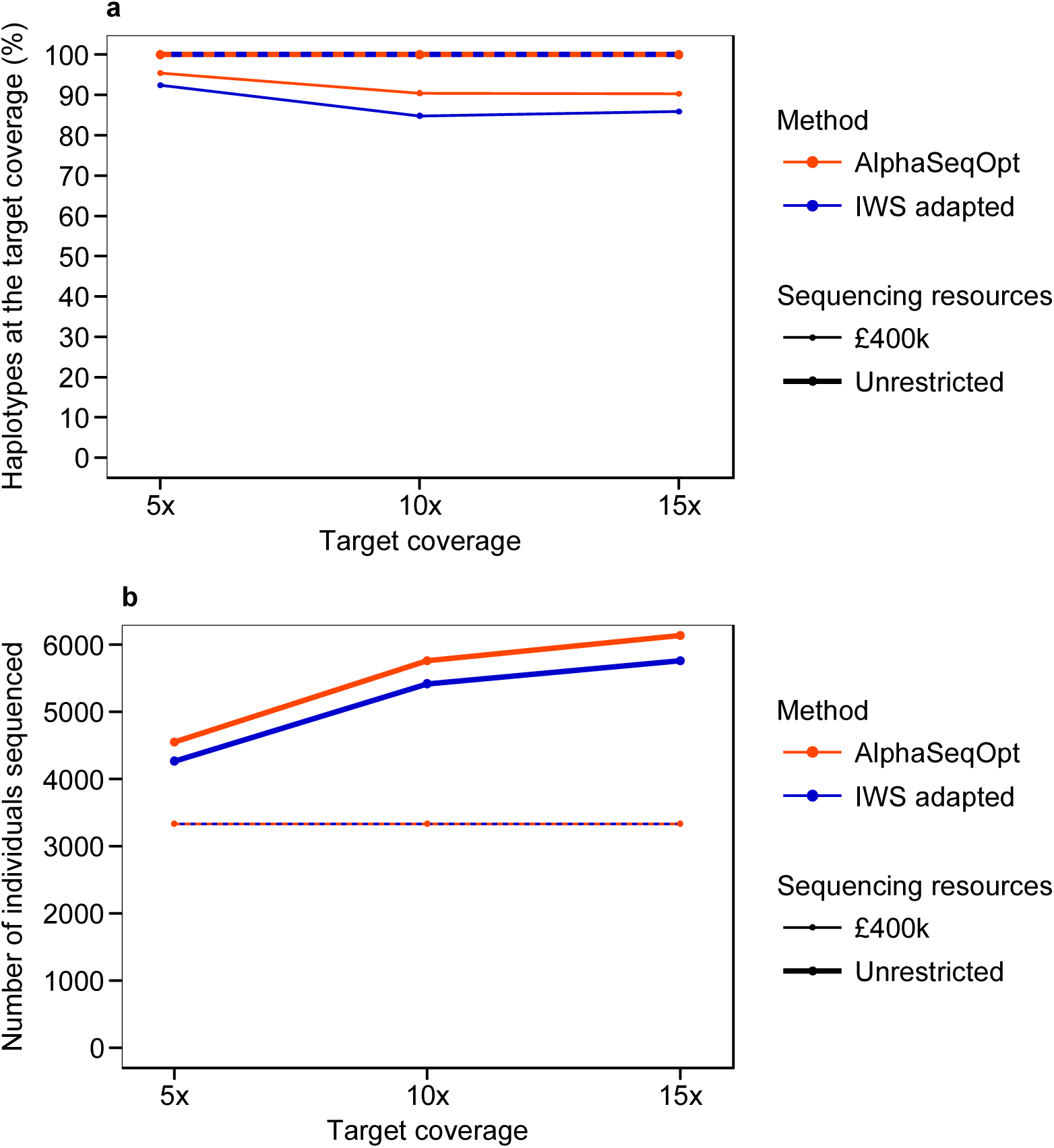
Performance of AlphaSeqOpt and the adapted IWS method when individual coverage is restricted to 1x, in terms of: (a) the percentage of population haplotypes sequenced at (or above) the target coverage; and (b) the number of individuals sequenced. When the target coverage was 5x, 10x, or 15x only haplotypes with population count ≥10, ≥20, or ≥30 were targeted, respectively.

With a budget of £400,000 a total of 3,333 individuals could be sequenced at 1x. Under this setting, AlphaSeqOpt delivered greater percentages of haplotypes sequenced at the target coverage than the adapted IWS method. If the budget was unrestricted, IWS selected a smaller set than AlphaSeqOpt to sequence all the targeted haplotypes at the desired coverage.

#### Effect of the number of exchanges per round during refinement

For the refinement of the set, there was an optimum number of exchanges per round that maximized the percentage of haplotypes sequenced at the target coverage given a fixed total number of exchanges. Figure 7a shows the percentage of haplotypes sequenced at (or above) the target coverage with a fixed number of total exchanges but with different numbers of rounds and exchanges per round, considering two extreme costs of library preparation. Figure 7b shows the size of the resultant set.

**Figure 7.**
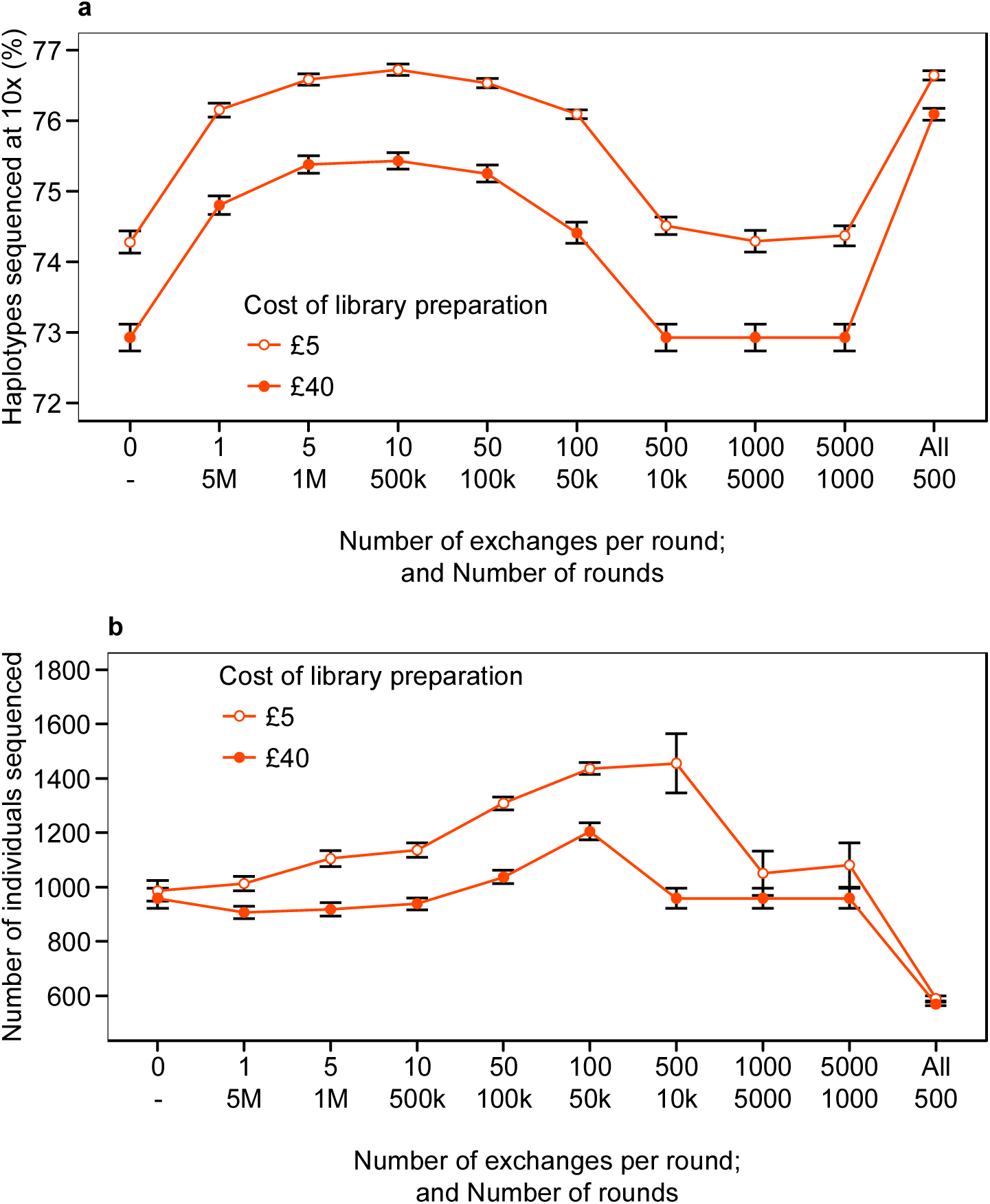
Effect of the number of exchanges per round: (a) on the percentage of haplotypes sequenced at (or above) the target coverage of 10x; and (b) on the number of individuals sequenced, with two costs of library preparation. The total number of exchanges is 5 millions in all cases. The sequencing resources were set to £800,000 and haplotypes with population count ≤2 were excluded from the analyses.

Doing 1 to 100 exchanges per round improved the percentage of haplotypes sequenced at the target coverage of the refined set to similar values. In this case, the set that produced the maximum percentage of haplotypes sequenced at the target coverage was obtained by doing 10 exchanges per round. Even though this greater percentage was generally achieved by increasing the number of unique sequenced individuals, the size of the refined set slightly decreased when the library cost was high and few exchanges per round were made. Doing more than 500 exchanges per round did not improve the results of the initial set when library cost was £40 and made the algorithm less robust when library cost was £5. However, the most extreme scenario of exchanging the whole set, which is equivalent to selecting a new initial set without any refinement in each round, provided the best improvement of the initial percentage of haplotypes sequenced at the target coverage and the greatest reduction of the sequencing set.

#### Filtering of rare haplotypes based on flanking context

As the target haplotype coverage increases, the least frequent haplotypes can be sequenced at the target coverage only if a large amount of resources is available. More sequencing resources can be focused on sequencing common haplotypes if the number of rare haplotypes included in the analyses is reduced either by excluding them all or by filtering them based on their flanking context.

Figure 8 shows the distribution of the sequencing resources depending on the population count of the haplotypes with the three different approaches to deal with the rare haplotypes: to include all singletons and doubletons in the analysis, to exclude them, or to filter them based on their flanking context. Almost half of the haplotypes in the test population were observed only once (singletons; 31.7%) or twice (doubletons; 13.2%), making a total of 3,971 singletons and doubletons. Of these, 953 (19%) remained after filtering based on their flanking context and these were considered as likely to have derived from a recombination event of two common haplotypes. We only show results for the scenarios in which the target haplotype coverage was 10x, with the total available sequencing resources being £400,000, £800,000, or £1,600,000.

**Figure 8.**
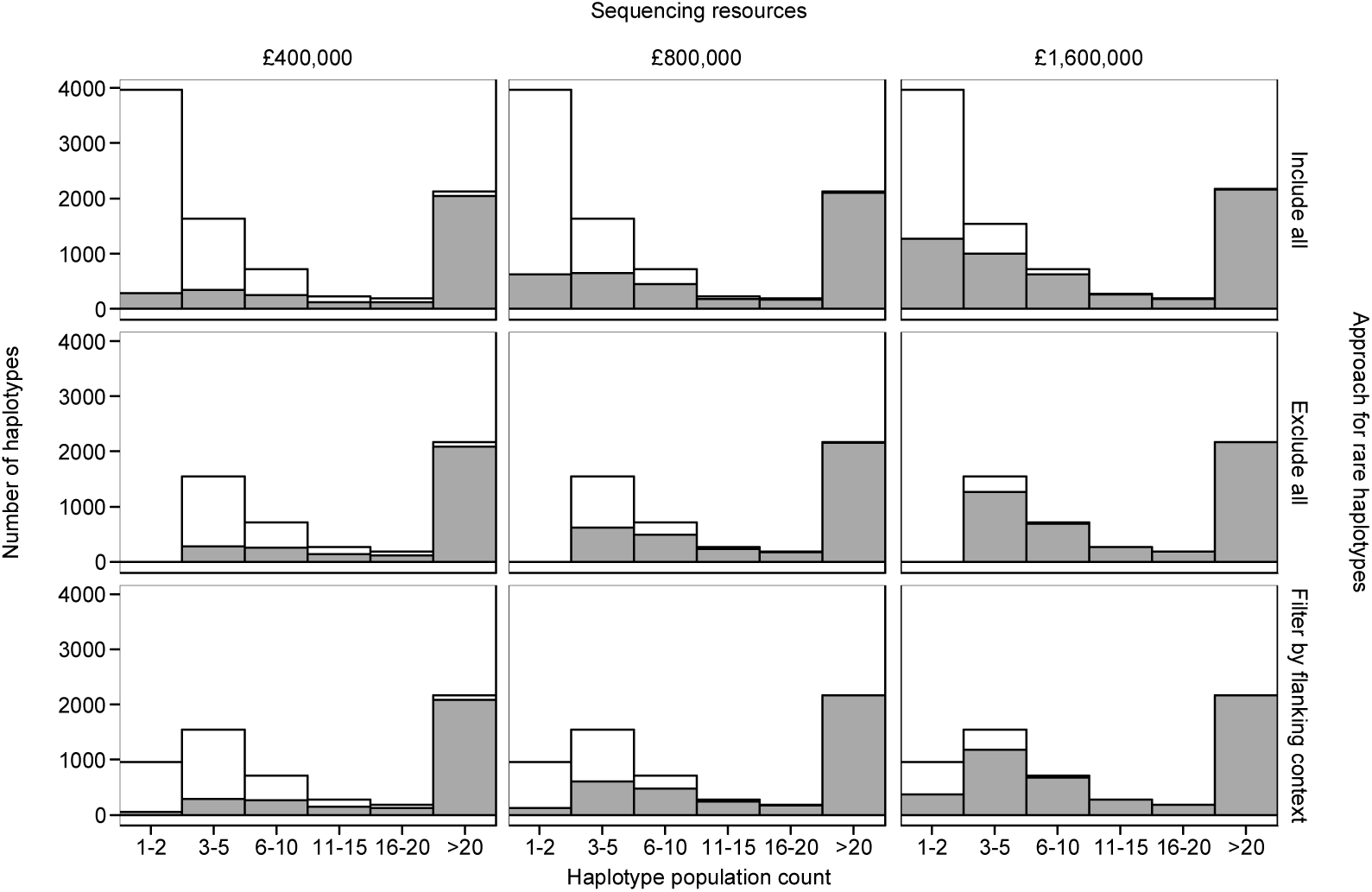
Number of haplotypes sequenced at (or above) the target coverage of 10x (filled section) using three different approaches to handle the rare haplotypes: to include all singletons and doubletons, to exclude all singletons and doubletons, or to filter them based on flanking context. Numbers are shown by haplotype population count and for different amounts of sequencing resources. The number of singletons and doubletons for each approach were 3,971, 0, and 953, respectively.

With £800,000, when all singletons and doubletons were included in the analyses 72.4% of the haplotypes with population count ≥3 were sequenced at (or above) 10x. This percentage increased to 75.3% when all singletons and doubletons were excluded. This percentage also increased, but a little bit less, when they were filtered based on their flanking context (74.8%). A similar trend was observed with £400,000 and £1,600,000.

When we have a large amount of sequencing resources we may be interested in targeting rare haplotypes as well as common haplotypes. By filtering based on their flanking context we can target the rare haplotypes that are likely to derive from a recombination of common haplotypes. With £1,600,000, a total of 38.6% of the 953 target singletons and doubletons were sequenced at 10x. Only 33.0% of these 953 was sequenced at 10x when all singletons and doubletons were included in the analyses without any restriction. This benefit of filtering by flanking context was not observed when less sequencing resources were available, probably because in such scenarios sequencing resources were implicitly focused on the common haplotypes.

## Discussion

We have presented a method that identifies which individuals need to be sequenced and at what coverage they should be sequenced when a given amount of sequencing resources are available so that the maximum percentage of the haplotypes present in the population are sequenced at (or above) a coverage that is sufficiently high to ensure that the consensus haplotypes can be accurately derived. Deriving the consensus haplotypes accurately is a critical requirement for achieving high population-based imputation accuracy under the LCSeq strategy and we have observed with a prototype of a novel population-based phasing and imputation method that there is a relationship between the coverage that a particular haplotype accumulates across individuals and the imputation accuracy of the consensus haplotype (Additional file 1: Figure S1). We also developed and tested a new approach to deal with rare haplotypes by filtering them based on their flanking context rather than excluding them from the analysis. We compared AlphaSeqOpt with previously published methods and hereafter discuss the advantages and limitations of AlphaSeqOpt.

### Advantages of AlphaSeqOpt over other methods

AlphaSeqOpt has two features that make it effective for its purpose: (1) a score function based on the frequency of the haplotypes in the sequencing set relative to the target coverage instead of on the population frequency of the haplotypes; and (2) a step of refinement of the initial set.

#### Score function

The score function that we propose allocates sequencing resources such that the percentage of haplotypes sequenced at any target coverage is greater than with other score functions based on the population frequency of the haplotype. The score function based on the population frequency of the haplotype used in the IWS method [17] was designed for producing the least redundant set that should be sequenced to have all the targeted haplotypes sequenced. The reduction of redundancy with the IWS method is achieved by giving a greater score to the least frequent haplotypes and, therefore, selecting the individuals that carry less frequent haplotypes first. Therefore, if the sequencing resources are sufficient for sequencing all the targeted haplotypes, the IWS method does so by sequencing a smaller set than AlphaSeqOpt. However, the IWS method is not ideal for identifying the set of individuals that would provide a more even sequencing coverage of the largest percentage of population haplotypes when the sequencing resources are limited and insufficient for sequencing all the targeted haplotypes at the desired coverage. With a score function that uses the population frequency the haplotype scores are constant until these haplotypes reach the target coverage, at which point they are set to zero. A score function based on the frequency of the haplotypes in the sequencing set relative to the target coverage like the one used in AlphaSeqOpt performs better for this purpose because, in contrast, the haplotype scores change as the sequencing resources are allocated. With the AlphaSeqOpt score function all haplotypes start with an equal score of 1 and their score increases exponentially as they approach the target coverage.

By doing this, the AlphaSeqOpt score function prioritizes the haplotypes that are already closer to the target coverage and, implicitly, the individuals that carry a larger number of these haplotypes. This reduces the percentage of haplotypes that are sequenced at a suboptimal coverage, but it increases the percentage of haplotypes that receive no coverage at all. With limited sequencing resources, AlphaSeqOpt selects a set for sequencing with a larger percentage of population haplotypes at the target coverage than the IWS method. These sequencing sets can be even smaller than the ones produced with IWS score function if the initial set is not refined.

#### Refinement of the initial set

The other main feature of AlphaSeqOpt is the step of refinement of the initial set. The step of refinement adjusts the allocation of resources by replacing individuals that have become redundant after the last additions to the set or by reducing the sequencing coverage of these individuals. A refinement step as described here further increases the percentage of haplotypes sequenced at the target coverage obtained with the AlphaSeqOpt score function. A side benefit in the context of LCSeq is that the refinement step achieves this increase by diversifying the set of individuals that are sequenced. While the IWS score function restrains the number of sequenced individuals, the AlphaSeqOpt score function benefits from low library costs relative to the total amount of sequencing resources available to produce larger sets with more unique individuals that are sequenced at lower coverage. This benefit is greater when the cost of library preparation represents a small fraction of the total amount of sequencing resources for LCSeq. Methods for reducing significantly the costs of library preparation for high-throughput LCSeq have already been described [27]. Increasing the number of individuals sequenced would empower subsequent imputation for more individuals (i.e., these individuals and their relatives) as well as any downstream analyses [12].

The refinement step can be fine-tuned by adjusting parameters such as the number of exchange rounds and the number of exchanges per round. The optimal parameters may depend largely on the size and structure of each dataset, but the following general observations were made:

- AlphaSeqOpt was very robust across repetitions. A stable solution was produced after a relatively low number of exchange rounds (unpublished results). Small further increases of the percentage of haplotypes sequenced at the target coverage could be obtained by using a longer chain of exchange rounds, but the benefit of this was little.

- To some extent, increasing the number of exchanges per round enables greater mobility across possible sets. Consequently, the algorithm can retrieve a better solution more easily. However, when too many exchanges are made per round, the benefit of this refinement of the existing set is diluted due to the drift towards solutions that are too divergent from each other and thus the final solution becomes less reliable. Exchanging all the individuals in the set is an extreme case of this that is equivalent to choosing the best of multiple initial sets without refinement. It can produce good results in terms of percentage of haplotypes for small sequencing sets.

#### Practical implications for real populations

Provided that the cost of library preparation is low enough or by restricting the maximum coverage of the individuals, AlphaSeqOpt will produce large sets of individuals with many unique individuals that are sequenced at low coverage.

The performance of AlphaSeqOpt will likely be influenced by structure of the data, either intrinsic, like the number and size of the chromosomes in a species or the degree of relatedness between individuals, or extrinsic, like the core length used to define the haplotypes. AlphaSeqOpt assumes that coverage is uniform along the genome but variation in coverage at the level of nucleobase should be expected, as well as variation of coverage between samples.

Although the criterion that is maximised in AlphaSeqOpt is the percentage of unique haplotypes sequenced at (or above) the target coverage, the method also provides good coverage in terms of total population haplotypes, i.e., haplotypes weighted by their population frequencies. Implicitly, the scores of more frequent haplotypes will increase faster than the scores of less frequent haplotypes because they are more likely to be carried by the individuals that are added to the sequencing set. In all scenarios tested, both AlphaSeqOpt and the adapted IWS method provided total percentages of haplotypes >99%, but the percentage was consistently greater for AlphaSeqOpt. Although both methods were similarly successful in covering the haplotypes of most of the population, AlphaSeqOpt captured a greater diversity of haplotypes at the desired coverage.

The resolution of the haplotype library will depend on the density of the marker array used to construct it. However after sequencing the individuals it is possible that haplotypes that were considered to form a single consensus haplotype when defined with marker data actually correspond to a number of true haplotypes. In such cases the sequence data can be clustered into the multiple consensus haplotypes and the pedigree information could enhance their imputation.

### Utility of filtering rare haplotypes based on flanking context

We proposed an approach that uses the haplotype population frequencies at the cores flanking a particular core to identify those rare haplotypes that could have derived from a recombination event. Although rare haplotypes may contain relevant biological information, we may not be able to impute and estimate accurately the effect of most rare haplotypes. The rationale behind the filtering approach that we propose is that sequence data of those rare haplotypes that are potentially mosaic of common haplotypes could enable a more precise capture of the recombination events that have occurred in the population and that this sequence data would also contribute to the consensus haplotypes of the haplotypes that gave rise to the mosaic. The new approach that we propose, although not ideal, may be of a particular interest in cases in which large amounts of sequencing resources are available.

In real populations we expect to identify large numbers of rare haplotypes. Preliminary tests indicated that in real populations our filtering approach based on flanking context can filter out around 92% of the singletons and doubletons observed, with the other 8% retained as potentially mosaic (unpublished results).

The challenge of targeting rare mosaic haplotypes is that the individuals that carry them must be sequenced at a greater coverage so that the rare haplotypes reach the target coverage. Another approach, for which we do not show results here, involves setting a lower secondary target coverage for less frequent haplotypes. This is a compromise solution where reducing the sequencing coverage of the rare haplotypes will reduce their imputation accuracy but will allow more rare haplotypes to be sequenced. In the particular case of potentially mosaic rare haplotypes having less coverage would be less critical because the information of the common haplotypes from which they derive will be also available. Any of the approaches discussed to filter low-frequency haplotypes can be combined using multiple frequency thresholds.

### Suitability of AlphaSeqOpt for low-coverage sequencing designs

A number of optimisation methods that use haplotypes derived from existing phased marker array genotypes have already been proposed to identify which individuals should be sequenced under the KeySires approach. Druet et al. [5] proposed a method that maximizes the proportion of haplotypes observed in the population that are sequenced. This method was more effective in detecting rare variants (minor allele frequency <5%) than methods based solely on pedigree information and it provided good imputation accuracies for both common and rare variants. Bickhart et al. [17] proposed the IWS method, which reduces the number of individuals that need to be sequenced in order to have all haplotypes above a certain population frequency sequenced. The method by Gusev et al. [16] selects the individuals that share the largest proportion of the population haplotypes with other individuals identical-by-descent (IBD). More recently, Neuditschko et al. [18] proposed a method based on the eigenvalue decomposition of a genomic relationship matrix that identifies individuals that maximise genetic diversity within complex population structures. These four methods identify which individuals should be sequenced but do not make any decision on the coverage at which they should be sequenced. Gonen et al. [14] proposed an approach that distributes sequence at variable coverage across individuals in a population. This method accounts for haplotype frequency and the ability to phase the resulting sequence data as criteria when choosing individuals to sequence and assigning sequencing coverage to those individuals and their recent ancestors. We have presented a method for optimizing the allocation of sequencing resources for the LCSeq approach so that the imputation accuracy of consensus haplotypes is high enough for enabling novel population-based imputation methods.

In practice, it is likely that a combination of the three sequencing approaches discussed in this paper (KeySires, LCSeq, and VarCoverage) would yield similar or even better imputation accuracies than LCSeq alone [28]. AlphaSeqOpt is flexible in that it can take into account the already available sequence information. Therefore, AlphaSeqOpt for optimizing LCSeq can be used either alone or complementarily to other existing methods to top-up the coverage of those haplotypes that are under-sequenced after using any other method. In this later case, however, existing methods or newly developed ones should be integrated to find the right allocation of resources into each of the three sequencing approaches.

## Conclusion

We have presented a method for optimizing the allocation of sequencing resources so that the maximum proportion of population haplotypes are sequenced at a coverage that is sufficiently high for population-based imputation with low-coverage sequencing. The haplotype score function and the refinement step make AlphaSeqOpt more effective for this purpose than methods reported previously for reducing sequencing redundancy. AlphaSeqOpt can account for sequence information already available for the population, which makes it a good complementary method to increase coverage of haplotypes that are not sufficiently covered when other optimisation methods are used. We also explored a new approach to deal with rare haplotypes by targeting only those that are likely derived by recombination of common haplotypes. This approach frees resources to sequence greater proportion of distinct haplotypes, which can be useful particularly when large amounts of sequencing resources are available.

## Authors’ contributions

JMH and RRF designed the algorithm and the study; RRF performed the analyses; RRF wrote the first draft; GG and SG assisted in the interpretation of the results and provided comments on the manuscript. All authors read and approved the final manuscript.

## Acknowledgements

The authors acknowledge the financial support from the BBSRC ISPG to The Roslin Institute BB/J004235/1, from Genus PLC and from grant numbers BB/M009254/1, BB/L020726/1, BB/N004736/1, BB/N004728/1, BB/L020467/1, BB/N006178/1 and Medical Research Council (MRC) grant number MR/M000370/1. This work has made use of the resources provided by the Edinburgh Compute and Data Facility (ECDF) (http://www.ecdf.ed.ac.uk). The authors thank Dr Andrew Derrington (Scotland, UK) for assistance in refining the manuscript.

## Additional files

**Additional file 1: Figure S1**

Format: pdf

Title: Expected haplotype imputation accuracy against the accumulated haplotype sequencing coverage, as estimated using a novel population-based imputation method (Battagin and Hickey, unpublished).

Description: A description of the prototype algorithm developed for the imputation of consensus haplotypes under the LCSeq approach and the simulated results on which the AlphaSeqOpt method is based. We generated 1x sequence data for the sires from a simulated population. The x-axis represents the expected accumulated coverage that each haplotype would receive. The y-axis represents the percentage of alleles phased and imputed for each haplotype. The imputation accuracy increased with the accumulated haplotype coverage until it platooned. Haplotypes with a sequencing coverage of 10x accumulated from 20 individuals sequenced at 1x were imputed to the whole population with an accuracy of 0.88. Haplotypes with a sequencing coverage of 15x or 20x accumulated from 30 or 40 individuals sequenced at 1x were imputed to the whole population with an accuracy of 0.93 or 0.97, respectively. For accurate inference of a consensus haplotype a certain amount of sequencing coverage must be accumulated. According to the results above, 10x or 15x could be good target coverages for the haplotypes.

